# Robust residue-level error detection in cryo-electron microscopy models

**DOI:** 10.1101/2022.09.12.507680

**Authors:** Gabriella Reggiano, Daniel Farrell, Frank DiMaio

## Abstract

Building accurate protein models into moderate resolution (3-5Å) cryo-electron microscopy (cryo-EM) maps is challenging and error-prone. While the majority of solved cryo-EM structures are at these resolutions, there are few model validation metrics that can precisely evaluate the local quality of atomic models built into these maps. We have developed MEDIC (Model Error Detection in Cryo-EM), a robust statistical model to identify residue-level errors in protein structures built into cryo-EM maps. Trained on a set of errors from obsoleted protein structures, our model draws off two major sources of information to predict errors: the local agreement of model and map compared to expected, and how “native-like” the neighborhood around a residue looks, as predicted by a deep learning model. MEDIC is validated on a set of 28 structures that were subsequently solved to higher-resolutions, where our model identifies the differences between low- and high-resolution structures with 68% precision and 60% recall. We additionally use this model to rebuild 12 deposited structures, fixing 2 sequence registration errors, 51 areas with improper secondary structure, 51 incorrect loops, and 16 incorrect carbonyls, showing the value of this approach to guide model building.

## INTRODUCTION

While technological advances in cryo-electron microscopy (cryo-EM) have made it possible to resolve protein complexes to resolutions rivaling X-ray crystallography [1], protein heterogeneity has limited the resolution for the majority of complexes, with 78% of cryo-EM maps deposited in the past year reporting a resolution worse than 3Å [2]. As the resolution drops from 3 to 5Å, modeling becomes increasingly difficult; the carbonyls become indistinguishable from the backbone density, side chain details are lost, and eventually, the backbone trace is no longer visible. Hand-built models at these resolutions can contain sequence registration errors, poor secondary structure, improper tracing of the backbone through the density, and incorrectly placed backbone carbonyls. There are several instances of models that have been deposited and published with errors that are found later by the community [3,4]. Methods like AlphaFold and RoseTTAFold [5,6] may help in alleviating these errors, but these methods’ inability to model structures with multiple conformations and their limited accuracy in modeling protein complexes will still lead to model errors.

Previous efforts in identification of model errors rely on metrics that primarily fall into one of two categories: model quality metrics that focus on atomic geometry [7,8], and fit-to-density metrics that focus on local fit-to-density [9-13]. Model quality metrics, such as the fraction of Ramachandran outliers, are not precise enough to catch local mistakes at these resolutions. Refinement protocols can easily push a wrong model to have good quality under these metrics. CaBLAM addresses this by defining a dihedral for the carbonyls in relation to the backbone trace and identifies when this angle deviates from expected values; however, due to its high cutoff value, CaBLAM is unsuitable for residue-level accuracy [11]. Density-based metrics have two major weaknesses: many are noisy at the level of individual residues and are better suited to evaluate a model’s global quality [12,13], while density-based metrics that robustly evaluate local fit rely heavily on side chain density, making them less reliable at resolutions below 3.5Å [14]. Furthermore, microscopists have a tendency to overfit their models to low-resolution density, so density fit by itself is not always enough to evaluate whether an error has been made [15].

Here, we present MEDIC (Model Error Detection in Cryo-EM), a statistical model that weighs the contributions of structural information with local model-map agreement to identify residue-level errors in a cryo-EM structure. The structural features of our model include both energy-guided metrics and a predicted error from a machine learning model trained to discriminate native and decoy structures. The use of a machine learning model to assess model geometry allows evaluation of non-bonded interactions such as hydrophobic burial, making it robust when used with lower-resolution data. We combine these structural features with a measure assessing agreement to density conditioned on data collected at a wide range of resolutions. We show reliable detection of errors on a set of 28 structures which were later solved to higher resolutions. On a smaller set of 12 deposited structures, we correct over 100 mistakes marked by our protocol with existing tools. Finally, we demonstrate that MEDIC can guide rebuilding in areas where AlphaFold models cannot.

## RESULTS

An overview of training and usage of MEDIC is shown schematically in Figure 1. MEDIC is trained to predict a probability of error for every residue, based on three features (Figure 1A): energy guided metrics for Ramachandran angles and bond deviations from Rosetta’s energy function [16], expected fit-to-density for a residue given the local resolution and the amino acid identity, and predicted model error from DeepAccuracyNet [17].

**Figure 1.**
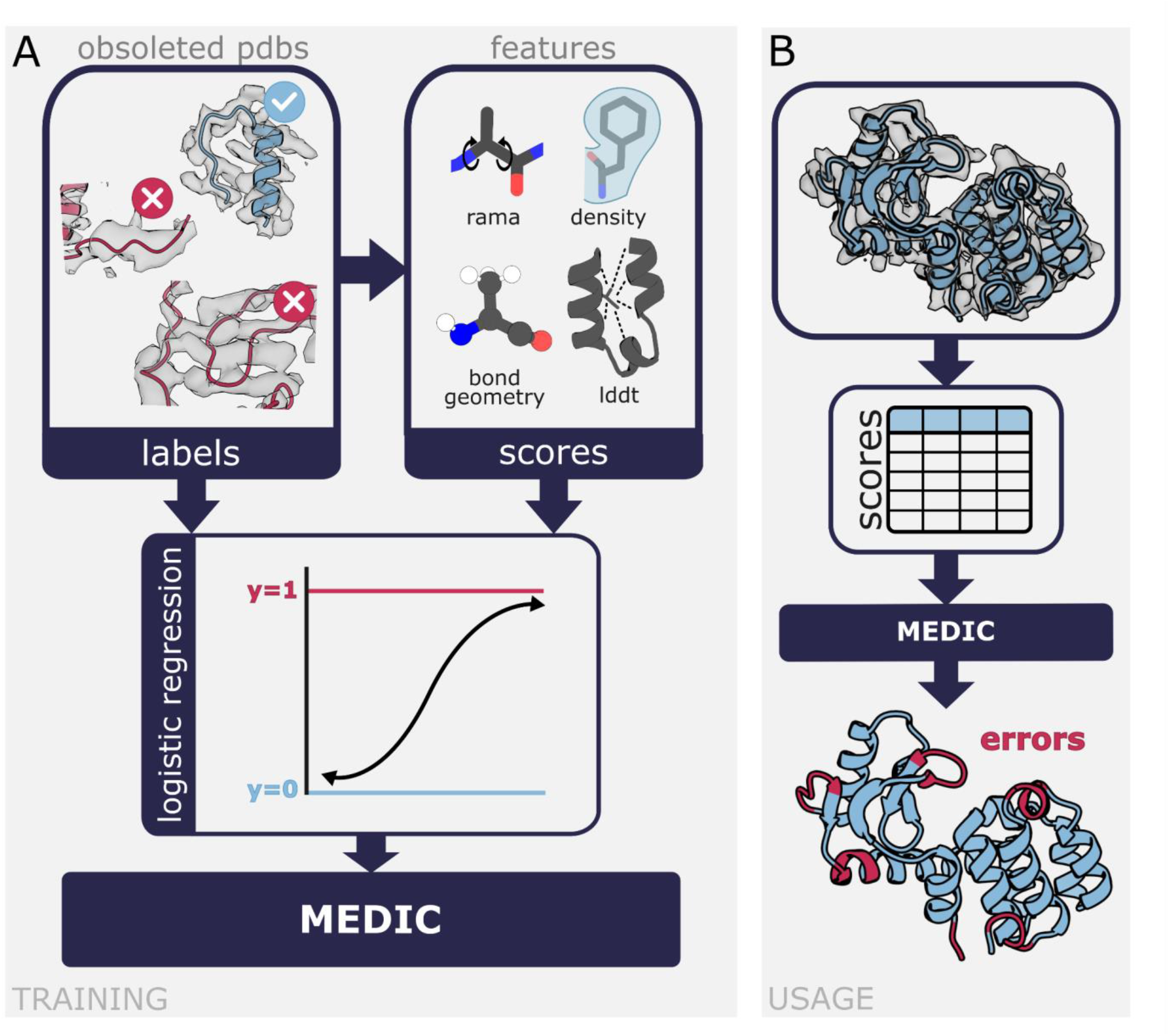
Overview of training and usage of MEDIC. **(A)** Pulling pdbs that had been edited after deposition, we marked every residue for which the backbone moved between the two versions as an error (red) and collected scores from each of our features on all residues. These labels and scores were fed to logistic regression, which gives us the statistical model, MEDIC. **(B)** To use MEDIC, provide a map/model pair to the program. We calculate the scores for each of our features, which are then passed to MEDIC. MEDIC predicts a probability that each residue is an error, where higher probability is indicative of an error.

DeepAccuracyNet is a deep convolutional neural network trained to distinguish native protein structures from Rosetta-determined decoys. It predicts per-residue local Distance Difference Test (lDDT), a measure of the number of atom pair distances that are maintained between a native structure and a decoy [18]. For our fit-to-density metric, we used masked real-space cross-correlation to measure density fit, and then normalize that value based on statistics for each residue identity at its local resolution, gathered from a set of deposited map-model pairs between the resolutions of 1.5 to 5Å.

Given these three features, a combined model was trained using a set of seven obsoleted protein structures which had been edited months after the initial deposition, presumably to correct structure errors. Our combined logistic regression model was trained to predict the residues that changed between the original and most-recent deposition. We validated this initial model on an additional 3 obsoleted structures which had been withheld from training. We compared MEDIC’s error probabilities to the residues that changed between these depositions and found that MEDIC had a precision of 76% at a recall of 60% (Supplemental Fig 1). Given the high performance on this initial set, we then used this model to generally evaluate deposited structures (Figure 1B). Throughout our analysis, we divide these probabilities into three categories: definite error, possible error, and non-error (see Methods). Data analysis is performed only with the high probability errors, while each image is colored according to the three categories.

### Validation on low resolution structures later solved to higher resolutions

To validate our approach, we considered EMDB-deposited structures between 3.5 and 5Å resolution, which were subsequently solved to better than 3.5Å (and at least 1Å better than the original deposition). There were 68 cases, of which we manually removed 40 with domain orientation changes between the high-resolution and low-resolution structures. The results on this dataset are summarized in Figure 2A. On this set of 28 structures, our method has a precision of 68% at a recall of 60%. This compares favorably with the widely used density-only metric, Q-score [9], which has a precision of only 35% at the same 60% recall.

**Figure 2.**
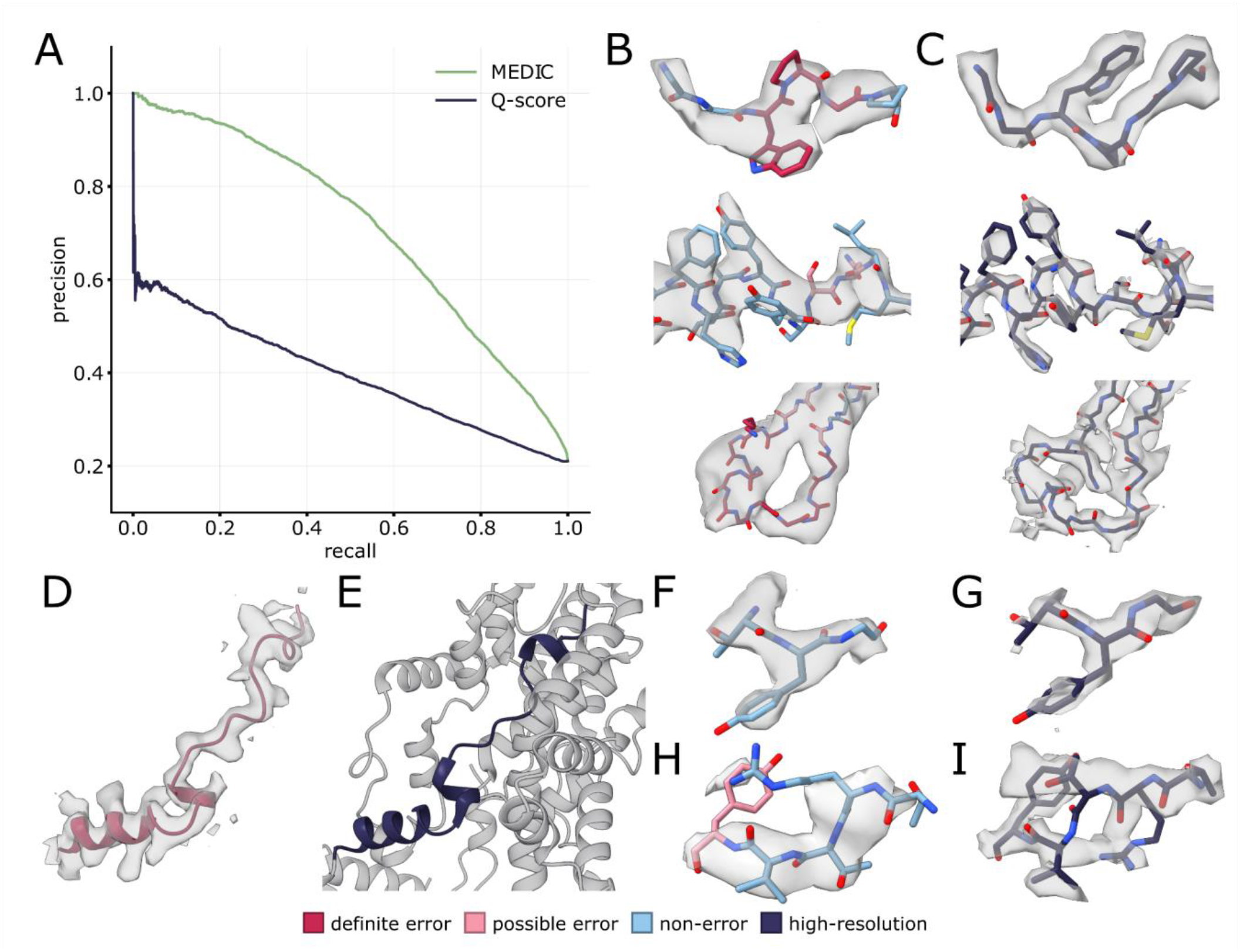
Results on validation set of analogous low-resolution and high-resolution structures. For panels B, D, F, and H, residues are colored by MEDIC error prediction. **(A)** Precision-recall curve of MEDIC error prediction and Q-scores on differences between low-resolution and high-resolution structures. **(B)** Examples of successful identification of errors in low-resolution structures: voltage-gated calcium channel (PDB 5GJW) **(top)**, insulin degradation enzyme (PDB 6B70) **(middle)**, transmembrane channel (PDB 6M66) **(bottom) (C)** The analogous region in the high-resolution structure: voltage-gated calcium channel (PDB 6JPA) **(top)**, insulin degradation enzyme (PDB 7K1F) **(middle)**, transmembrane channel (PDB 6WBF) **(bottom). (D)** False positive predicted by MEDIC in ATP synthase (PDB 6F36). **(E)** High-resolution structure ATP synthase (PDB 6RD5) with missing context from low-resolution structure colored in gray. **(F)** MEDIC misses an incorrect carbonyl in low-resolution structure of a dehydrogenase (PDB 7E5Z). **(G)** Analogous region in high-resolution structure (PDB 7VW6). **(H)** MEDIC does not mark a region in the dehydrogenase (PDB 7E5Z) that matches the high-resolution data. **(I)** Mistake made in the high-resolution model (PDB 7VW6).

We next examined which features were predictive of the true positives identified by MEDIC. Approximately 81% are predicted by the lDDT alone, while the remaining 19% require at least 2 features to be considered an error. The reliance on lDDT to predict most of the errors could be because of bias in the training set, which primarily contains long segments that were corrected. It might also simply reflect the types of errors microscopists tend to make; hand-built models are much more likely to fit the density well but have poor geometry and structural features.

Some of the errors identified by MEDIC in the low-resolution structures are highlighted in Figure 2B, with the corresponding model in its high-resolution density map in Figure 2C. In a structure of a voltage-gated calcium channel (PDB 5GJW), it is difficult to trace the backbone while properly accounting for the large aromatic side chain density (Figure 2B, top panel). The mistake is identified by MEDIC with relatively equal contributions from the lDDT and bond geometry scores. Likewise, the error found in an insulin degradation enzyme (PDB 6B70) is captured by multiple features, this time the density and bond geometry scores (Figure 2B, middle panel). The backbone is hardly visible in the density map, which may explain why the microscopists had difficulty properly fitting the serines into the density. In contrast, the mistake found in a transmembrane channel (PDB 6M66) is dominated by the lDDT score (Figure 2B, bottom panel). It would be difficult to catch this error by visual inspection, as the model seems reasonable given the density.

To better understand any shortcomings of MEDIC, we looked at two structures for which our performance was worse than the aggregate results. In a partial complex of an ATP synthase (PDB 6F36), MEDIC falsely marks an entire stretch of residues as a mistake (Figure 2D) because it does not see the proper structural context for this particular sequence as it is unmodelled in the low-resolution structure (Figure 2E). The other case which MEDIC performed poorly on, a dehydrogenase (PDB 7E5Z), contained many errors fewer than 3 residues in length which MEDIC failed to identify, two of which are shown in Figures 2F-I. We fail to mark an incorrect carbonyl as an error in the low-resolution model (Figure 2F) that is supported by the higher-resolution data (Figure 2G). However, we find zero high-probability errors in a region of the low-resolution model (Figure 2H) which appears to be an error in the high-resolution model (Figure 2I).

Given our worse performance on the errors in the dehydrogenase (PDB 7E5Z), we manually examined 30 differences across 4 low-resolution structures that MEDIC failed to identify. Among these, 16 were mistakes in the model built against low-resolution data, while 14 were either ambiguous in the high-resolution density or seemingly incorrect in the high-resolution model. Three examples are highlighted in Supplemental Figure 2: one difference where the high-resolution structure has an error (Supplemental Fig. 2A-B), and two more where the high-resolution structure is not supported by the density (Supplemental Fig. 2C-F).

### Using MEDIC to guide model rebuilding

With the understanding that MEDIC is relatively precise when identifying errors, we next wanted to assess the usefulness of the model to aid in a manual structure rebuilding process. To that end, we evaluated MEDIC on a selection of 12 models with diverse topologies and resolutions and attempted to fix – using Rosetta refinement tools and AlphaFold – all the segments marked as errors (see Methods). There were 237 segments predicted to be definite errors (with high error probability), 33 of which were disordered regions with little or no visible density (Supplemental Fig. 3). Of the remaining 204 segments, 133 (65%) were 1-3 residues in length, 38 (19%) between 4-9 residues, and 33 (16%) were greater than 10 residues. We were able to rebuild and fix 120 (59%) of these segments; for an additional 26 segments, we were able to significantly reduce the number of definite errors in that region. The fixable mistakes included 2 sequence registration errors, where the sequence is shifted on the backbone relative to the correct placement, 51 incorrect loops, 51 cases of poor secondary structure, and 16 flipped carbonyls (Table 1).

**Table 1.** Summary of identified and corrected high-probability errors in 12 deposited models, excluding disordered regions.

| **PDB ID** | **Total Segments**<br>**Marked by MEDIC** | **Fixed** | **Improved** |
| --- | --- | --- | --- |
| 6JT1 | 7 | 2 | 1 |
| incorrect loop | 3 | 2 | 1 |
| 7R9U | 8 | 6 | 1 |
| poor secondary structure | 5 | 4 | 1 |
| incorrect loop | 3 | 2 | 0 |
| 7QFQ | 30 | 26 | 1 |
| poor secondary structure | 16 | 16 | 0 |
| incorrect loop | 9 | 8 | 1 |
| incorrect carbonyls | 2 | 2 | 0 |
| 3J9E | 4 | 3 | 1 |
| incorrect loop | 2 | 1 | 1 |
| incorrect carbonyls | 2 | 2 | 0 |
| 7S9D | 24 | 16 | 1 |
| poor secondary structure | 11 | 10 | 1 |
| incorrect loop | 7 | 6 | 0 |
| 5MM4 | 56 | 25 | 9 |
| poor secondary structure | 17 | 14 | 3 |
| incorrect loop | 17 | 9 | 5 |
| incorrect carbonyls | 3 | 2 | 1 |
| 6C14 | 10 | 5 | 0 |
| poor secondary structure | 4 | 3 | 0 |
| incorrect loop | 2 | 2 | 0 |
| 6XOW | 4 | 3 | 0 |
| incorrect loop | 3 | 3 | 0 |
| 7VOJ | 1 | 0 | 1 |
| incorrect loop | 1 | 0 | 1 |
| 6DMB | 36 | 22 | 10 |
| poor secondary structure | 3 | 1 | 2 |
| incorrect loop | 18 | 10 | 8 |
| sequence registry | 1 | 1 | 0 |
| incorrect carbonyls | 9 | 9 | 0 |
| 6EIO | 11 | 9 | 0 |
| poor secondary structure | 3 | 3 | 0 |
| incorrect loop | 5 | 5 | 0 |
| sequence registry | 1 | 1 | 0 |
| 6C0V | 13 | 4 | 1 |
| poor secondary structure | 1 | 0 | 1 |
| incorrect loop | 4 | 3 | 0 |
| incorrect carbonyls | 1 | 1 | 0 |
| Totals | 204 | 120 | 26 |
| poor secondary structure | 60 | 51 | 8 |
| incorrect loop | 74 | 51 | 17 |
| sequence registry | 2 | 2 | 0 |
| incorrect carbonyls | 17 | 16 | 1 |

**Figure 3.**
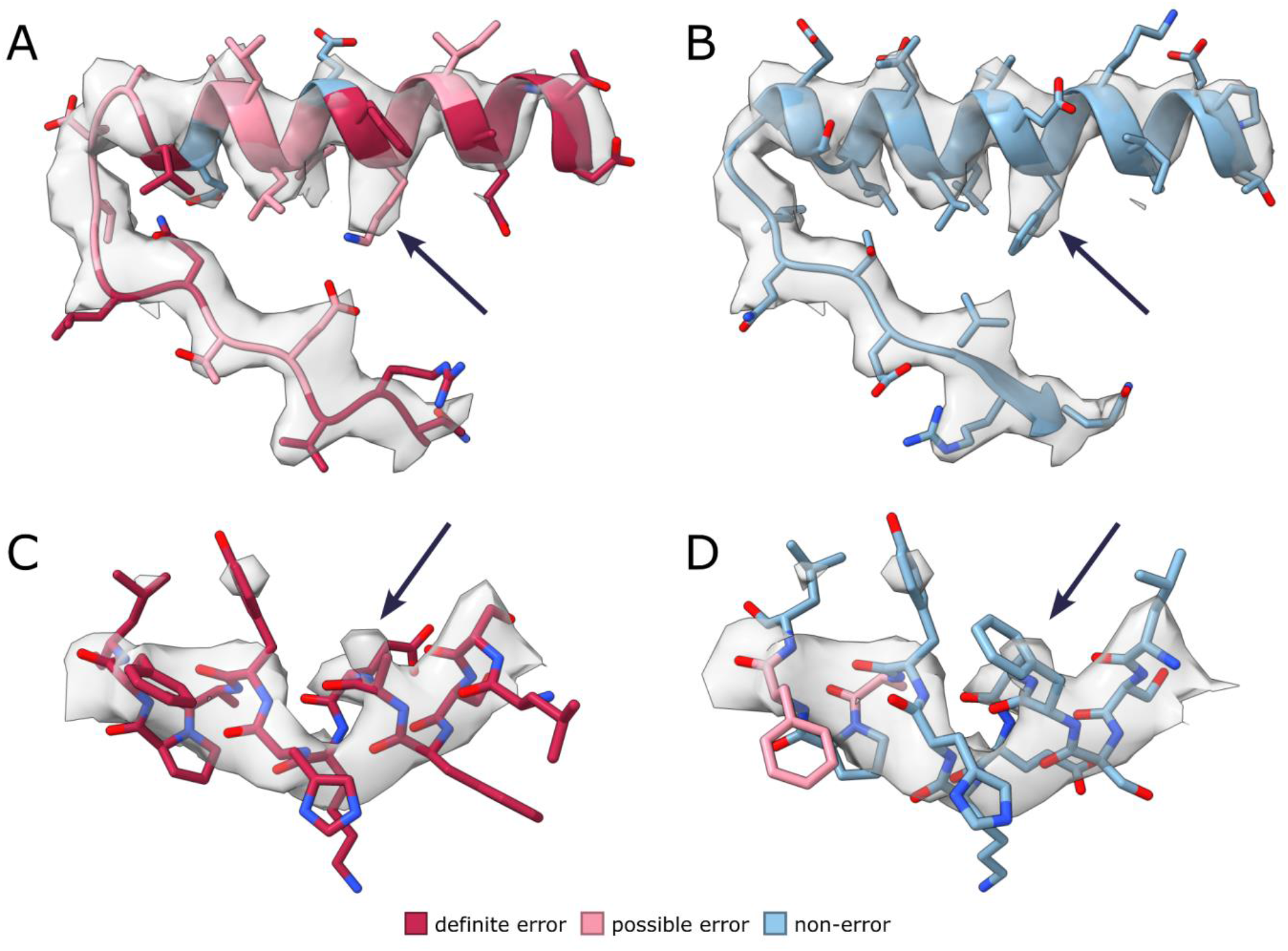
Sequence registration errors identified in deposited structures. All residues colored by predicted error from MEDIC. **(A)** Sequence registration error in lipid scramblase (PDB 6E1O). **(B)** Rebuilt model of **A**, where phenylalanine fills large side-chain density. **(C)** Sequence registration error in hedgehog receptor (PDB 6DMB). **(D)** Rebuilt model of **C** with a bulge added, where phenylalanine fills large side-chain density.

A representative subset of errors that our method was able to address are highlighted in Figures 3 and 4. In these cases, we were able to correct 2 significant sequence registration errors (Figure 3). Figure 3A compares the deposited structure of a lipid scramblase (PDB 6E1O) with our new model. Notably, our model has better hydrophobic packing and we explain the large side chain density with a phenylalanine as opposed to a lysine residue (Figure 3B). This sequence registration error was propagated from a previously solved crystal structure (PDB 4WIS), in which the density for this region was poorly resolved. In both structures, this helix is preceded and followed by unresolved regions, making proper sequence placement more difficult. Conversely, the sequence registration error found in a hedgehog receptor (PDB 6DMB) occurs because the pitch of the helix is not visible in the density (Figure 3C). The addition of a bulge in the repaired model (Figure 3D), justified by the preceding proline, pushes a phenylalanine into large side chain density which was poorly explained by an alanine in the original model.

**Figure 4.**
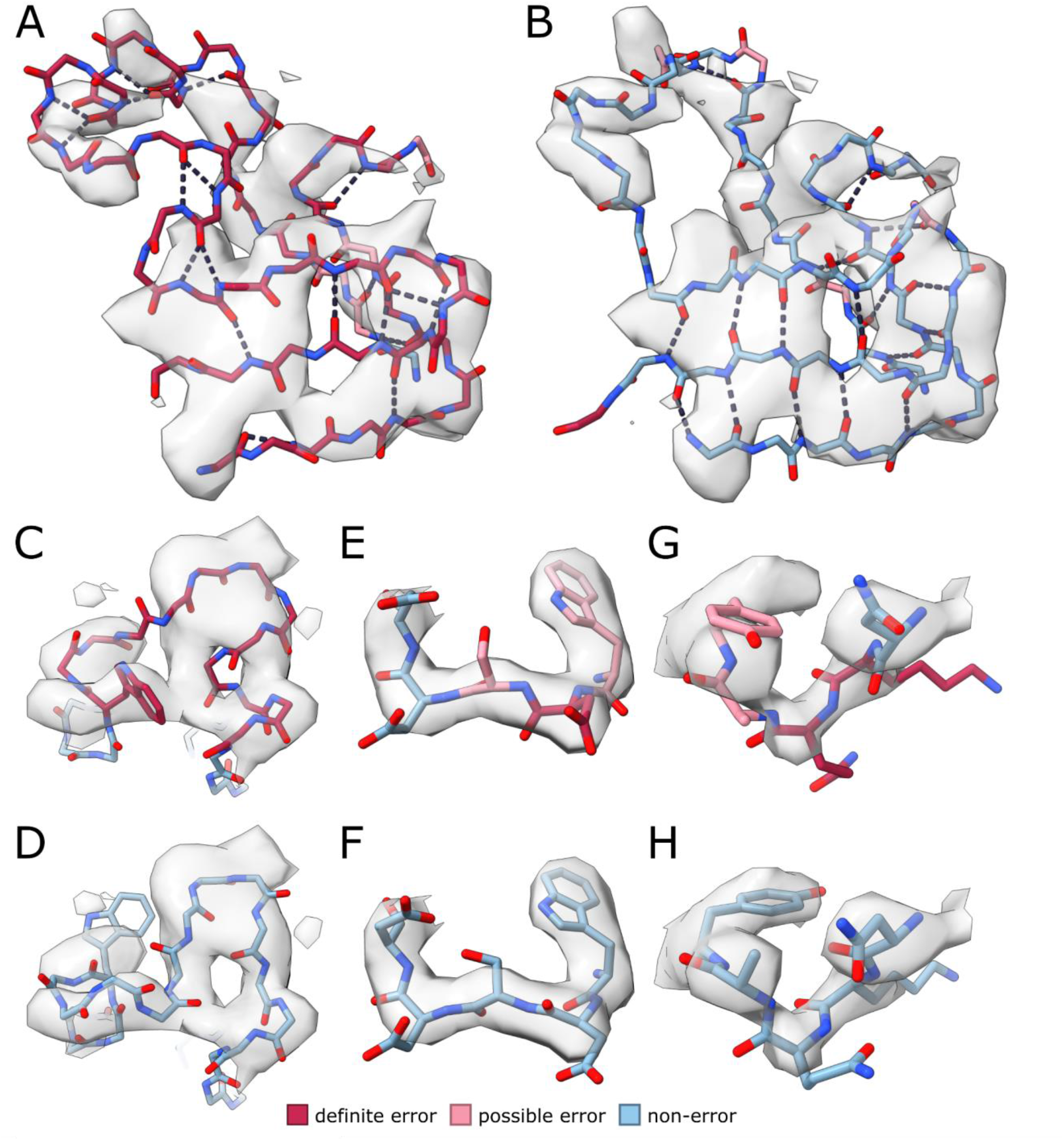
Backbone errors identified in deposited structures. All residues colored by predicted error from MEDIC. **(A)** Predicted errors in kinesin motor domain (PDB 5MM4) **(B)** Rebuilt model of **A** with better hydrogen-bonding. **(C)** Small loop in hedgehog receptor (PDB 6DMB) that poorly explains density. **(D)** Rebuilt loop of **C**, which has less unexplained density. **(E)** Protein backbone with incorrect carbonyls in bluetongue virus (PDB 3J9E). **(F)** Rebuilt backbone of **E** with improved Ramachandran angles. **(G)** Deposited structure in neurotoxin (PDB 7QFQ). **(H)** Rebuilt model of **G** with better fit to density and improved Ramachandran angles.

MEDIC is also capable of finding gross backbone errors, including areas with poor secondary structure and incorrect loops. In Figure 4A, it is clear by eye that the beta strands of this kinesin motor domain (PDB 5MM4) have poor hydrogen bonding. Upon fixing the secondary structure (Figure 4B), our method marks these regions as correct, as MEDIC balances proper structural features with density fit. In addition to identifying poor structural features, MEDIC can recognize if a stretch of residues is assigned the incorrect secondary structure, such as the region from a hedgehog receptor (PDB 6DMB) depicted in Figure 4C. However, our fixed model is supported by more than the lDDT score; it has less unexplained density, which is reflected by large improvements in the density scores (Figure 4D).

Furthermore, MEDIC can identify some shorter, subtler backbone errors, such as incorrectly placed carbonyls, by combining multiple features (Figure 4E-H). The deposited model of the bluetongue virus (PDB 3J9E) has a Ramachandran angle that falls just in the “Allowed” region (Figure 4E). MEDIC uses the lDDT and the bond geometry scores to predict this error, and after rebuilding, both Ramachandran angles and density fit improve (Figure 4F). Similarly, the structure for a neurotoxin (PDB 7QFQ) contains Ramachandran angles which Molprobity also classifies as “Allowed” (Figure 4G). We find this error with relatively equal contributions from lDDT, density, and geometry energies. The rebuilt model improves the density fit for the tryptophan and alanine residues while removing the problematic Ramachandran angles (Figure 4H). Of the over 1300 residues identified as errors across these 12 models, approximately 66.5% of them were predicted by the lDDT score alone, 1.4% by the density, and 0.4% by the Ramachandran energy, while 32% required at least 2 features.

To quantify MEDIC’s performance on this set of structures, we used the differences between the deposited structures and our rebuilt models (see Methods) to determine that MEDIC has a precision of 67% at recall of 60% (Supplemental Fig. 4). The increased performance of MEDIC at high recall values compared to the low-vs. high-resolution validation set could be attributed to a few factors. In the set of validation structures, it is possible that the high-resolution models may contain errors. Moreover, there could still be conformational differences between the high- and low-resolution structures, such as flexible loops or shifts that occur at interfaces contained in only one of the depositions. Both would hurt MEDIC’s perceived performance.

### Identifying errors in all deposited structures in the EMDB

After confirming MEDIC’s high accuracy and utility in model building, we ran MEDIC on all structures in the EMDB between the resolutions of 3 to 5Å to gauge the reliability of the method on over 1500 depositions. The aggregate statistics from this run are shown in Figure 5. Upon inspection, several models were composed of docked crystal structures with no visible density for one or more domains, so we removed residues with a model-map correlation of less than 0.4. In Figure 5A, we show the fraction of residues marked as errors in every EMDB deposition. There is only a slight trend with resolution, which is unsurprising given that as we move to lower resolutions, microscopists are more likely to dock crystal structures or use homology modeling than hand-build structures. Because cryo-EM maps are rarely homogenous in resolution, we also report the fraction of residues marked as errors after grouping by atomic B-factors (Figure 5B). At very low atomic B-factors (indicating well-resolved density), very few errors are made. As the atomic B-factors increase, more mistakes are made.

**Figure 5.**
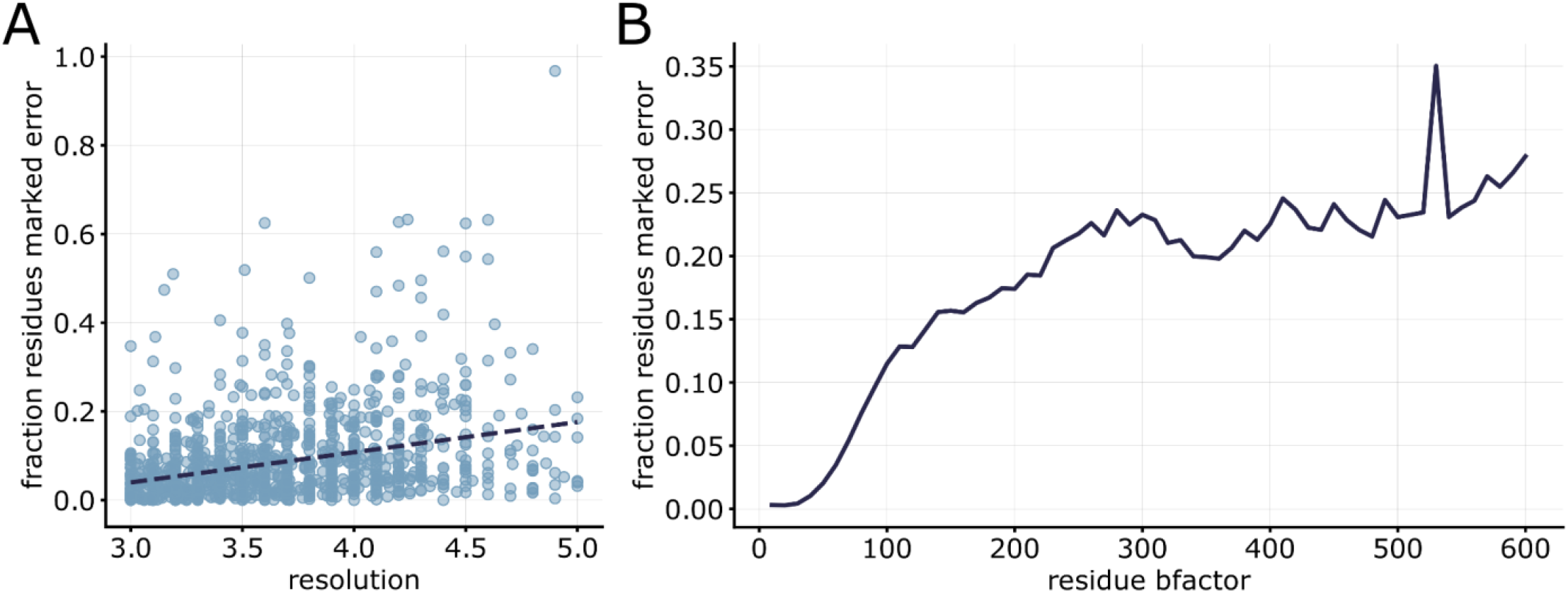
Aggregate statistics on all deposited structures in the EMDB. For both plots, residues with low density cross correlation, less than 0.4, were not included. **(A)** Fraction of residues marked as an error by MEDIC in each deposited structure. **(B)** Fraction of residues marked as an error with atomic B-factors between X-10 and X.

We manually inspected the outliers in the data: maps with very high error fractions, and errors with low atomic B-factors. Although the fraction of errors is greater than 40% on the 20 model-map pairs we examined, the errors do seem real. In some cases, entire domains have little to no secondary structure (Supplemental Fig. 5A-B). All these structures were built pre-Alphafold, using outdated (then state-of-the-art) structure prediction software or by hand-tracing into low-resolution data. Unsurprisingly, we find that 88% of the errors in this set are predicted by the lDDT alone. In the structures that contain errors with low atomic B-factors, we find that while some errors appear to be real, there also appear to be false positives. There are several causes for the perceived false positives, including residues marked as errors because they are involved with ligand or metal binding, or they correspond to very short disordered segments (Supplemental Fig 5C-E).

### Comparison to AlphaFold

Although it is clear that MEDIC can identify errors in hand-built structures, many microscopists will now start model-building from an AlphaFold prediction [19]. We compare MEDIC’s performance to AlphaFold models, highlighting loops which we identified as an error in the original deposition (Figure 6A & 6D) and where AlphaFold predictions do not fit the density. The loop predicted by AlphaFold for the motor protein, prestin (PDB 7S9D), would require significant rebuilding (Figure 6B). MEDIC identifies our new model, built with tools in Rosetta, as correct (Figure 6C). The shorter loop predicted by Alphafold for the bluetongue virus (PDB 3J9E) is not only a poor fit to density (Figure 6E); the carbonyls are placed incorrectly when compared to our final model (Figure 6F). Of the 12 models we rebuilt, 23 regions (from 7 different AlphaFold models) would have required rebuilding. AlphaFold was confident (predicted lDDT > 70) in 10 of these regions, which means that modelers would need to manually identify these mistakes, not just remove low confidence regions, and then rebuild, presumably by hand. MEDIC will be useful for this editing process: our method was able to identify that the deposited structure or our rebuilt model was correct in 18 of those 23 regions. In the remaining 5 cases, we were unable to build a structure that satisfied MEDIC.

**Figure 6.**
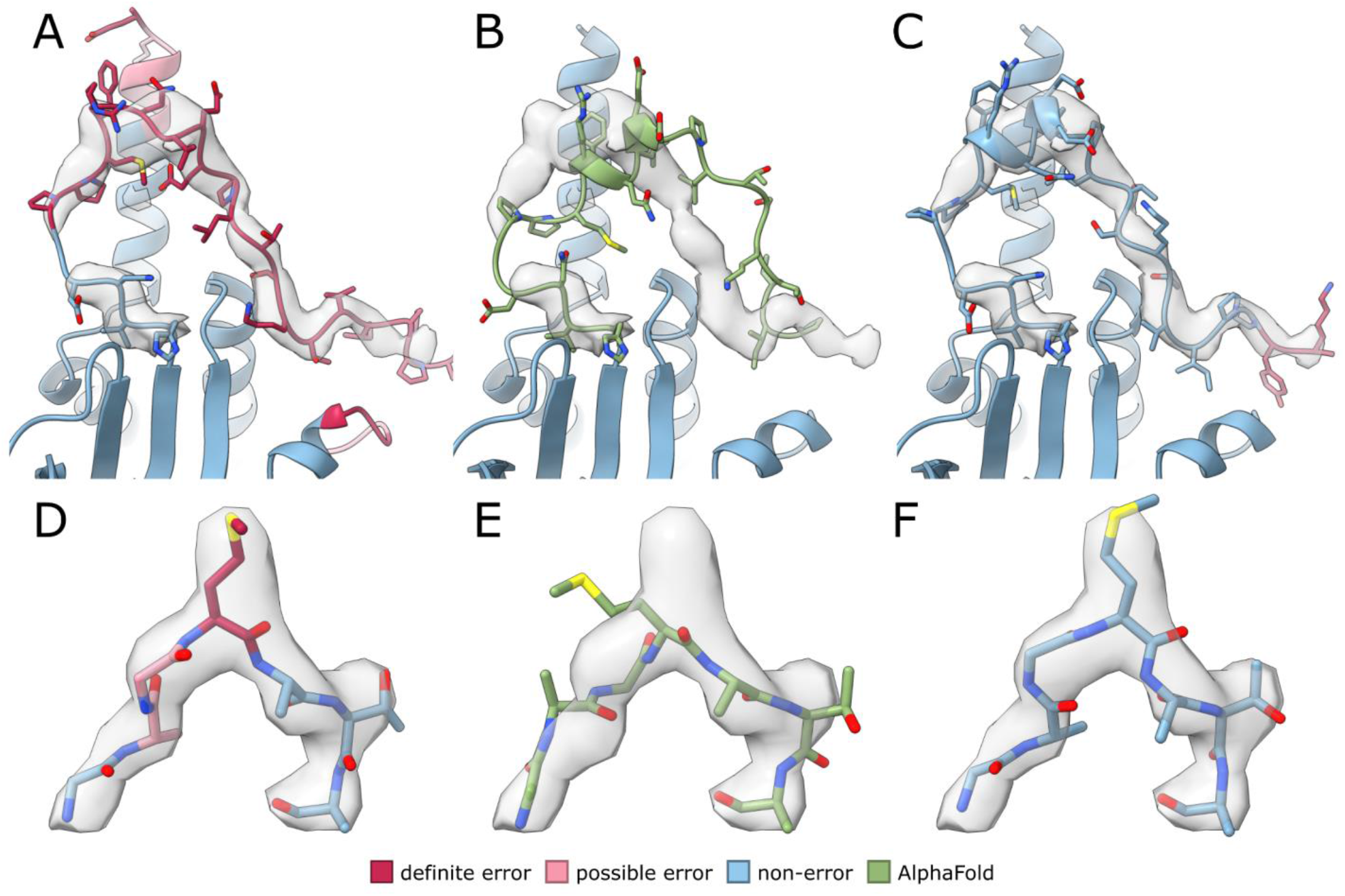
Errors identified by MEDIC where AlphaFold models do not match the density. **(A)** Deposited structure of prestin (PDB 7S9D), colored by error prediction. **(B)** AlphaFold model for prestin after docking the relevant domain into the density. **(C)** Rebuilt structure of loop in **C**, colored by error prediction. **(D)** Deposited structure for bluetongue virus (PDB 3J9E) in density map, colored by error prediction. **(E)** AlphaFold model for bluetongue virus after refining the model into the density. **(F)** Rebuilt structure of loop in **D**, colored by error prediction.

## DISCUSSION

In this report, we develop a method for the identification of local errors in cryo-EM models in the resolution range of 3-5Å. We validate our method on cryo-EM structures that have later been solved to higher resolutions and show that MEDIC has a precision rate 30% better than competing methods. We also highlight the use of MEDIC in the model building process by identifying and correcting over 100 errors in a set of 12 deposited models and demonstrating MEDIC’s use in conjunction with AlphaFold. While many of the errors are predicted by lDDT alone, we also find errors that make use of structure and density in tandem. Of the errors we examined, we noticed that MEDIC erroneously marks the following: prolines, termini, residues involved in binding, and regions where there is little to no supporting density (Supplemental Fig. 3). We believe prolines have a higher false positive rate because their geometry scores tend to be higher and caution users to be critical of isolated prolines which MEDIC calls errors.

As it becomes more commonplace to model large protein complexes into lower resolution density maps [20], validation metrics that can evaluate these structures and help guide rebuilding are necessary. MEDIC’s performance on structures with resolutions worse than 5Å has not been tested and given that our statistics for density did not include these resolutions, it is unclear how reliable our method will be in those cases. MEDIC could be extended to lower resolutions by gathering more statistics and by measuring density fit across longer stretches of sequence, making it suitable for use with cryo-electron tomography. A training set could be curated from low resolution structures which are later solved to higher resolutions by removing regions with different domain orientations and regions of ambiguity. Incorporating AlphaFold models into the training set may also be useful, so that MEDIC more explicitly learns to find regions which have good structural features but do not fit the density well.

AlphaFold has not only made it possible to model lower-resolution structures, it has drastically changed the model building process for higher resolution structures as well. Now microscopists will edit loops or interaction sites rather than build entire structures. For large complexes, identifying and fixing errors in AlphaFold models can still be error-prone and time consuming, especially if these are flexible regions solved to lower local resolutions. Creating a program to automatically dock these models and fix any remaining errors would reduce the amount of time and expertise necessary to solve structures. MEDIC could be used to guide this rebuilding process; our method’s high precision would substantially reduce the sampling space, which makes the problem of automatically fixing local errors much more tractable. Based on the observations described here, we believe that MEDIC will be a powerful validation tool for cryo-EM microscopists.

## METHODS

### Preparation of input pdbs

Preparation of pdbs for training or for error detection is a three-step process. First, we remove all ligands, nucleotides, or noncanonical amino acids. Then we refine the structure into the density map, first with cartesian minimization and then with Rosetta’s LocalRelax protocol [21]. Finally, we perform B-factor fitting on the refined model. After this, all the scores for the model features can be calculated.

### Structural features

The energy guided metrics in our model are pulled from Rosetta’s realistic energy function [16]. Every pdb is refined in Rosetta as described above, so that the energy scores are meaningful. Then, the energies for Ramachandran angles and bond deviations are evaluated for each residue in the structure and fed directly into the model.

The final structural feature, predicted lddt, comes from DeepAccuracyNet [17]. Because DeepAccuracyNet was trained on smaller structures, <300 residues in length, we run the model on portions of the structure at a time: a sequence of 20 residues and the context within 20Å of that query sequence. The predicted lDDT values are saved for only the query sequence and then passed to the model.

### Density feature

To calculate expected fit-to-density for amino acids, we collected statistics on a set of 24 deposited map-model pairs, using atomic B-factors as a substitute for local resolution. Each model and was prepared as described above. The masked real-space density cross-correlation was calculated for every residue and each was placed into a bin according to its amino acid identity and the average B-factor of the residues within an 8Å neighborhood. A mean of the cross-correlation scores was computed for each amino acid/B-factor bin and a standard deviation was calculated across each B-factor bin.

Now that we have collected statistics, we can apply them during error prediction. The means and standard deviations are used to transform the cross-correlation of each residue in a protein model into a z-score. A very negative density z-score is indicative of a residue which fits the density worse than expected, given its amino acid identity and the average B-factor. The density z-score is then passed to the model. This process of collecting statistics and transformation of raw scores is carried out for the cross-correlation of the residue by itself and the cross correlation of a three-residue window centered on the residue of interest.

### Training on obsoleted pdbs

We probed the RCSB for pdbs which had been edited after deposition, pulling all cryo-EM structures between 2.5 and 4Å resolution that had coordinates replaced [22]. Upon manual inspection, 10 models of the 46 were chosen, eliminating cases where changes were made to ligands, nucleotides or only rotamers, or where the obsoleted model didn’t resemble a globular protein. 3 of the 10 models were withheld from training and used for validation.

We now have a set of pdbs that contain mistakes made by microscopists and need to generate labels for training, marking each residue in a model as an “error” or “non-error.” We compare the obsoleted pdb with the newer version, removing any domains or regions that exist in only one of the models. Each residue in which the backbone atoms have an RMSD greater than or equal to 1Å between the two models is marked as an error. To capture sequence registration errors, any residue that appears in the obsoleted model but not the new version is marked as an error. This process resulted in approximately 800 errors out of a total of 21000 residues. We then trained a logistic regression classifier, with balanced class weights, to predict the errors using the structural and density features.

### Evaluation of error vs non-error

To determine the threshold at which a residue is an error, we chose a threshold value from the precision-recall curve which balances the two statistical measures. We use both the precision-recall from the 12 rebuilt models and the high-resolution low-resolution validation set to choose thresholds. We consider every residue with a probability above 0.78 to be a definite error. At threshold 0.78, MEDIC has a precision of 70% and recall of 80% on the set of 12 rebuilt structures and a precision of 78% and recall of 49% on the validation set. All statistics and data analysis are done only with this more stringent threshold value. We consider every residue with a probability between 0.78 and 0.6 to be a possible error. At a threshold of 0.60, MEDIC has a precision of 52% and recall of 95% on the 12 rebuilt structures and a precision of 68% and recall of 61% on the validation set. Every residue with a probability less than 0.6 is a non-error.

### Calculation of error contributions

To determine whether a single feature is predictive of an error, we take the probability equation that we have learned from the final training dataset (Eq. 1), where *l* is the lDDT score, *sd* the single residue density score, *ld* the 3-residue density score, *r* the Ramachandran energy, and *b* is bond energy:

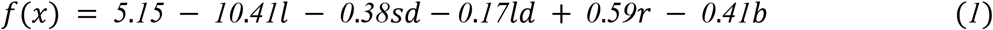

We replace all features, except the ones of interest, with the mean score, derived from the scores of the EMDB depositions (over 1500 cases). For example, we replace lDDT, Ramachandran and bond energies with the corresponding mean values to calculate how predictive the density scores are. We then take the result from Eq. 1 and plug it into Eq. 2 to get the final probability. If the final probability is above our threshold for definite errors, then that residue is predicted by a single feature.

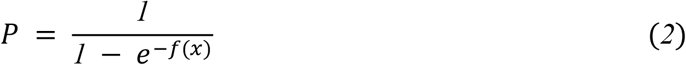

### Error identification on deposited structures and retraining

We identified cryo-EM structures with less than 2000 residues and a resolution between 3 and 5Å. We then chose 12 structures with diverse topologies and resolutions to run through our error detection, using the statistical model obtained from training on the obsoleted pdbs. We used a probability threshold of 0.62, derived from the precision-recall curve for the small set of 3 withheld obsoleted structures. We chose a slightly lower threshold, sacrificing precision (60%) for recall (85%) to ensure that we would find most of the errors.

After error identification, we attempted to rebuild every region that was predicted to be an error, following the protocol described below. We then added these 12 models to our training data. We generated error labels by looking first for residues with RMSDs greater than 1.5 after rebuilding, for which the probability was greater than 0.5 and had dropped by 0.2 after fixing.

We also labeled residues with RMSDs between 0.5 and 1.5 with probabilities greater than 0.6 and that dropped by 0.2. Any 1-residue errors from this set were removed if they were not within 2 residues of other errors. These labels and scores were passed into the logistic regression with the obsoleted pdbs, adding an additional 1200 errors to the dataset.

### Model rebuilding

For each rebuild, we ran AlphaFold on the sequence [5], docking the model or separately docking its domains into the density using UCSF Chimera [23]. Then, we removed all regions in the deposited model that were identified as errors plus/minus 2-3 residues on either side of the segment. We passed the AlphaFold models and the trimmed deposited model as templates to RosettaCM [24]. We ran at least 2 rounds of iterative RosettaCM, passing the top 5 models out of the total 50 to the next round. Additional rounds were run if model convergence for the top 5 was poor or if additional errors were detected by MEDIC and Molprobity. Any remaining regions which AlphaFold or RosettaCM were not able to fix were built with RosettaES [25]. Success in rebuilding was determined by how well regions matched the density by eye, Molprobity scores, and MEDIC predictions. All images of these structures were made in ChimeraX [26].

### High- and low-resolution structure validation

We pulled all cryo-EM structures between 3.5Å and 10Å for which there was another deposition with the same UniProt ID and at least 1Å higher resolution, with a maximum of 3.5Å. If the query structure had a model-map FSC greater than 10Å, the pair was thrown out. From this initial pool of 68 structures, 40 pairs were tossed because there were significant conformational changes caused by image processing, ligand binding, or physiological conditions.

For the remaining 28 pairs of structures, the high-resolution structure was docked and refined into the low-resolution map, and the low-resolution structure was refined into its own density [21]. The backbone RMSD between the two structures was calculated for every residue and all residues with at least 1Å RMSD were labeled as errors. Residues that only existed in one model of the pair were tossed and not used in validation. Error detection was then run on the low-resolution structure using the statistical model from the larger dataset and precision-recall curves were calculated with the described labels.

### Comparison to Q-scores

To obtain a precision-recall curve for Q-scores, we first generated Q-scores for each residue in the structure. We then subtracted the Q-score for the residue from the expected Q-score based on the global resolution for that map. This procedure mimics the usage of Q-score, where modelers are advised to examine residues which drop below the expected value. The difference between expected and actual Q-score is then used to calculate the precision-recall curve.

### Identifying errors in all deposited structures in the EMDB

We pulled every deposited cryo-EM structure with resolutions between 3 and 5Å, removing approximately 300 structures for which the model-map FSC at 0.5 was worse than 10Å. Then we prepared each pdb as described above and ran the statistical model from the combined dataset to detect errors. Of the 2037 structures that met our criteria, MEDIC successfully ran on 1713 (87.4%). To remove regions of disorder, we toss out all residues for which the density cross correlation is less than 0.4 in all subsequent analyses.

## Supporting information

EMDB supplemental data

## CODE AVAILABILITY

MEDIC will be made available for download at: https://github.com/gabriellareggiano/MEDIC

## ACKNOWLEDGEMENTS

Funding for this research was provided by NIH R01-GM123089 (FD, GR). We are grateful to Nao Hiranuma for DeepAccuracyNet support.

---

**Supplemental Figure 1.**
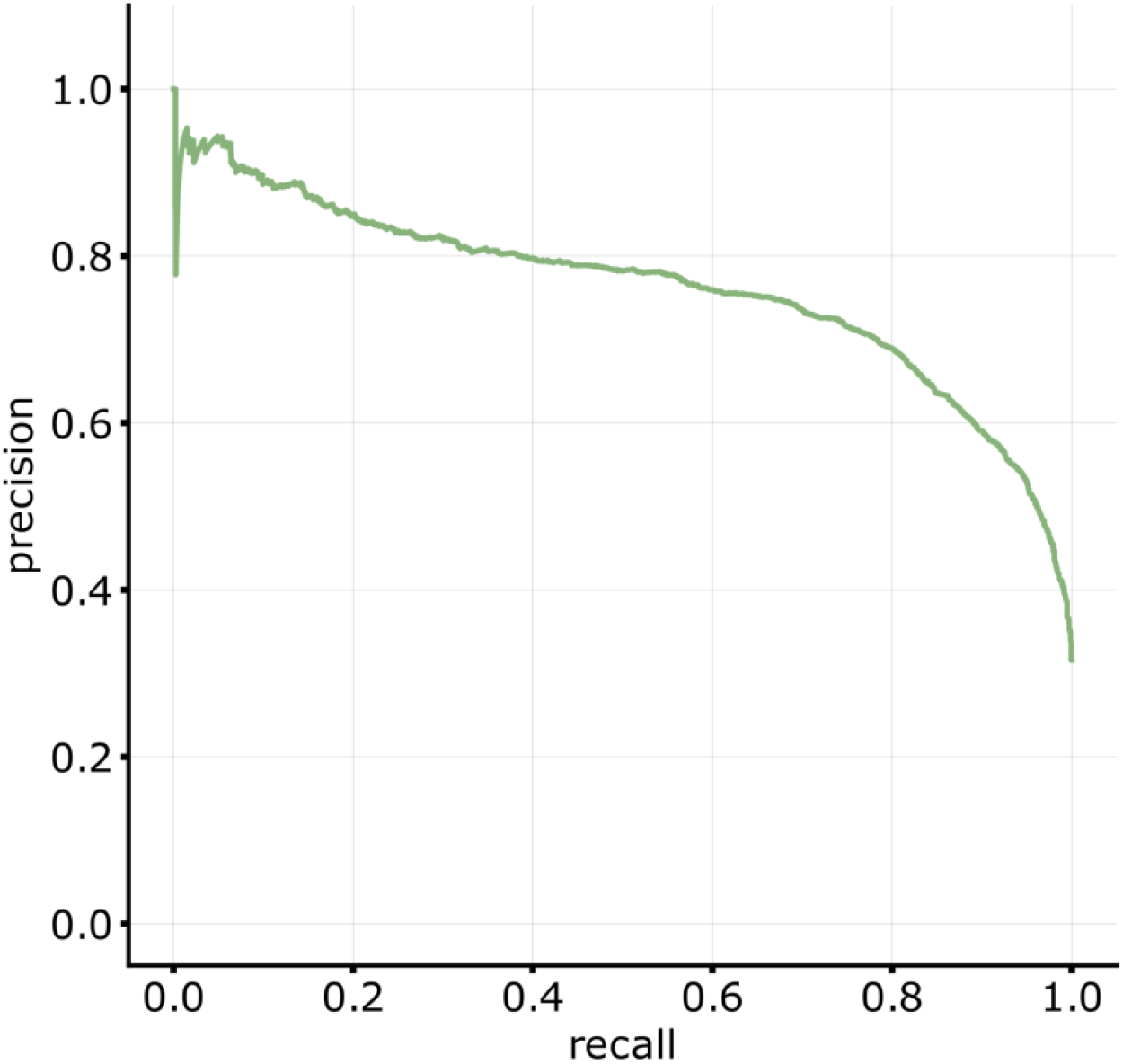
Precision-recall of MEDIC predictions for the set of 3 withheld obsoleted structures using an early version of MEDIC.

**Supplemental Figure 2.**
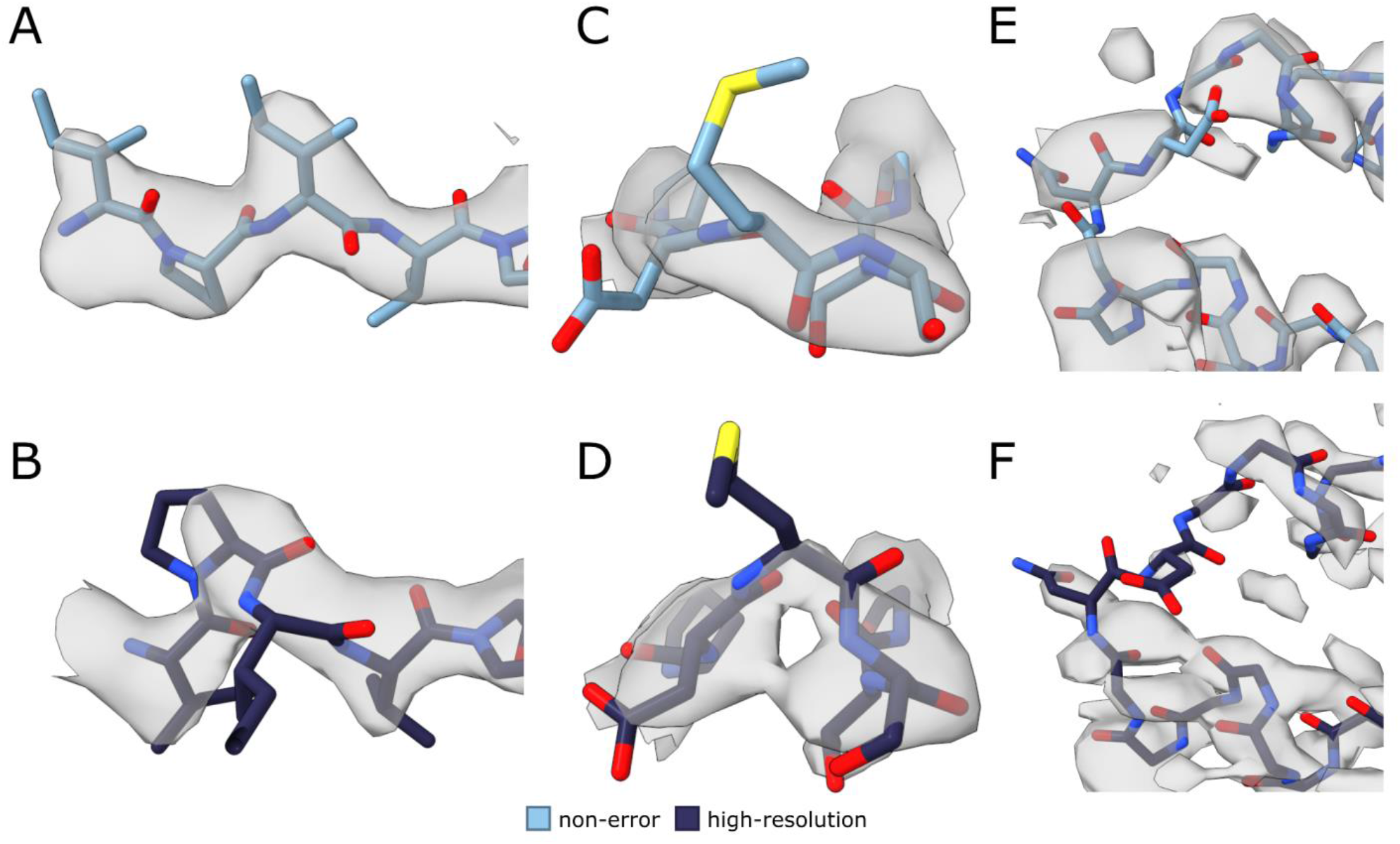
False negatives from the high-resolution vs. low-resolution validation set that are not supported by the high-resolution data. Low-resolution structures (A, C, E) colored by MEDIC prediction. **(A)** Low-resolution structure of glutamate hydrogenase (PDB 3JD3) that differs from high-resolution. **(B)** High-resolution structure of protein from **A** (PDB 5K12) appears to have an error in this region. **(C)** Loop in glutamate hydrogenase (PDB 3JD3). **(D)** High-resolution structure (PDB 5K12) is poorly resolved in the same region from **C. (E)** Region from low-resolution structure of TRPV5 (PDB 6PBE) is poorly resolved. **(F)** Corresponding high-resolution structure (PDB 7T6O) is poorly resolved in the same region from **E**.

**Supplemental Figure 3.**
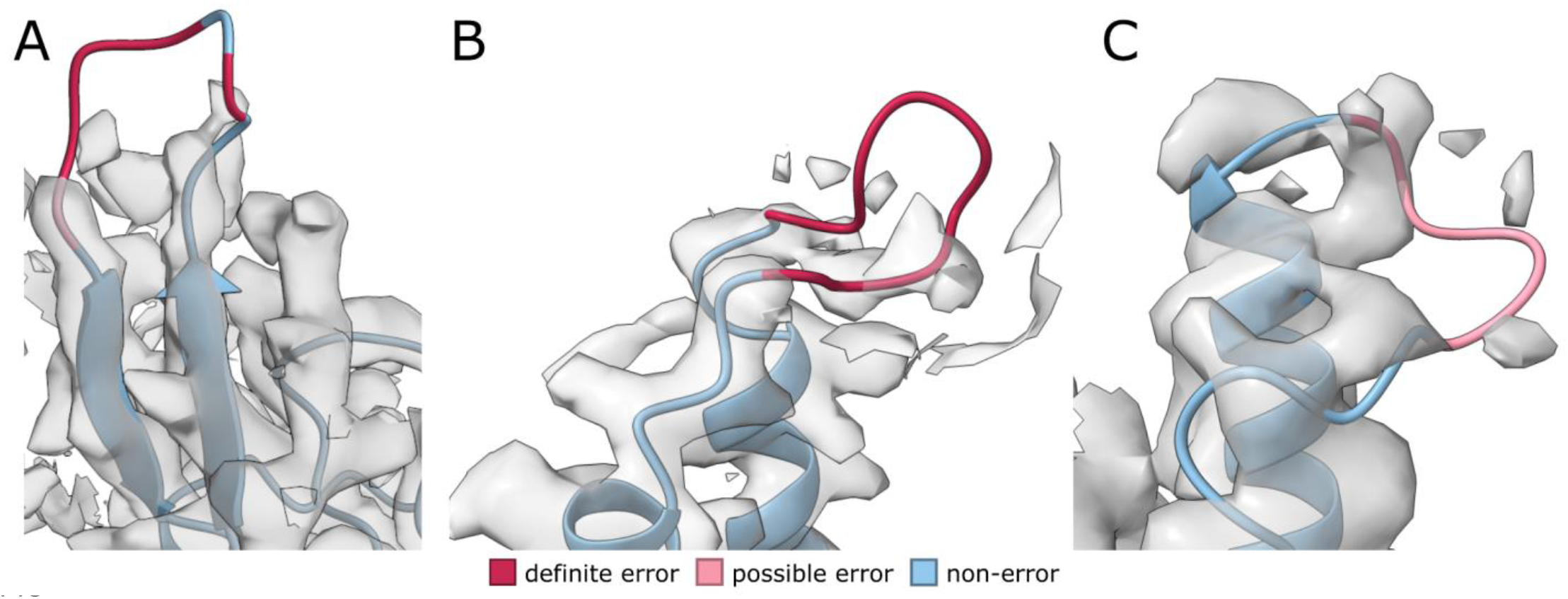
Regions with little supporting density marked as errors by MEDIC. Regions shown are from our new rebuilt models for the following structures: **(A)** neurotoxin (PDB 7QFQ), **(B)** lipid scramblase (PDB 6E1O), **(C)** prestin (PDB 7S9D).

**Supplemental Figure 4.**
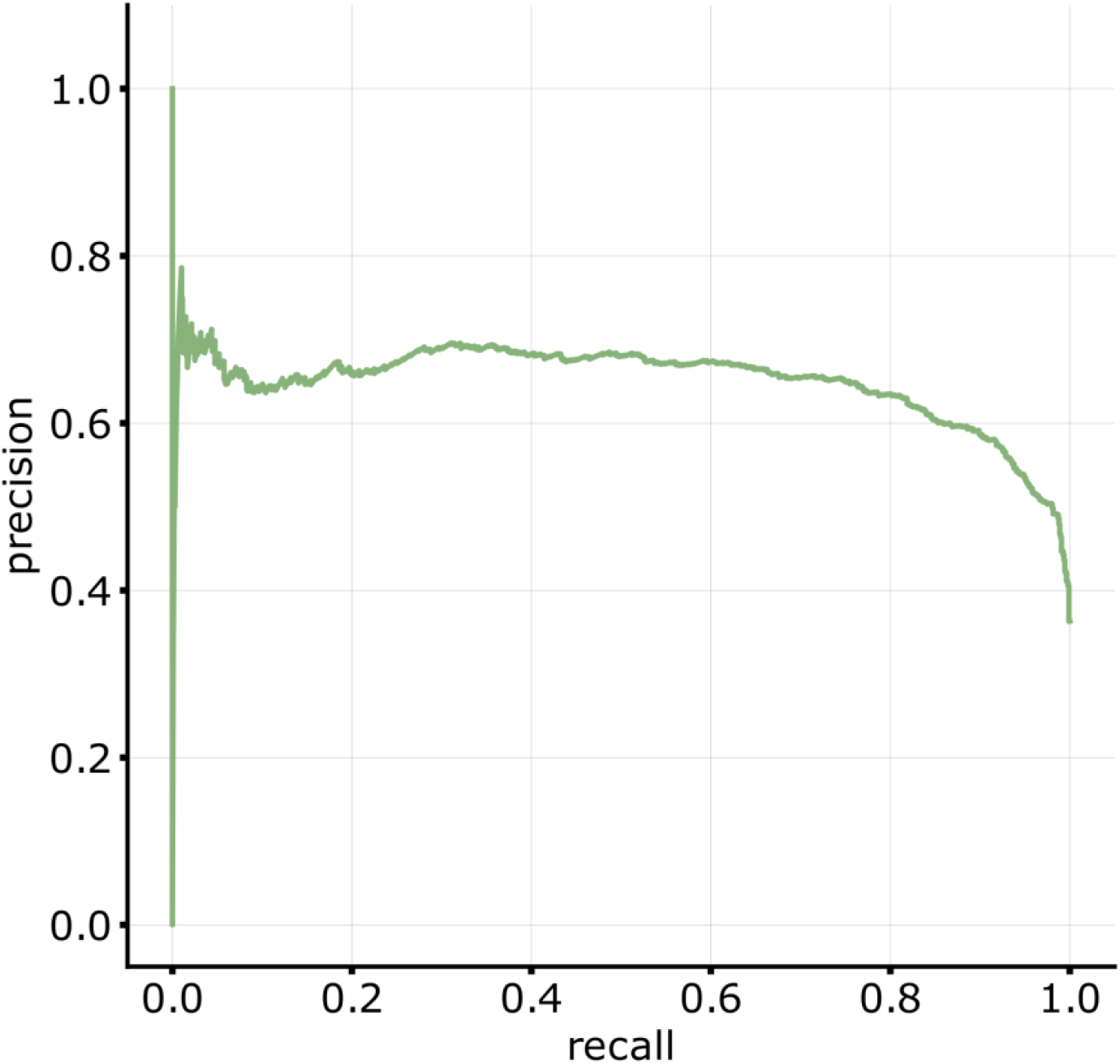
Precision-recall of MEDIC predictions for the set of 12 rebuilt structures using the full training dataset. We used leave-one-out validation on each of the models from the set of 12 rebuilt structures to avoid bias in training.

**Supplemental Figure 5.**
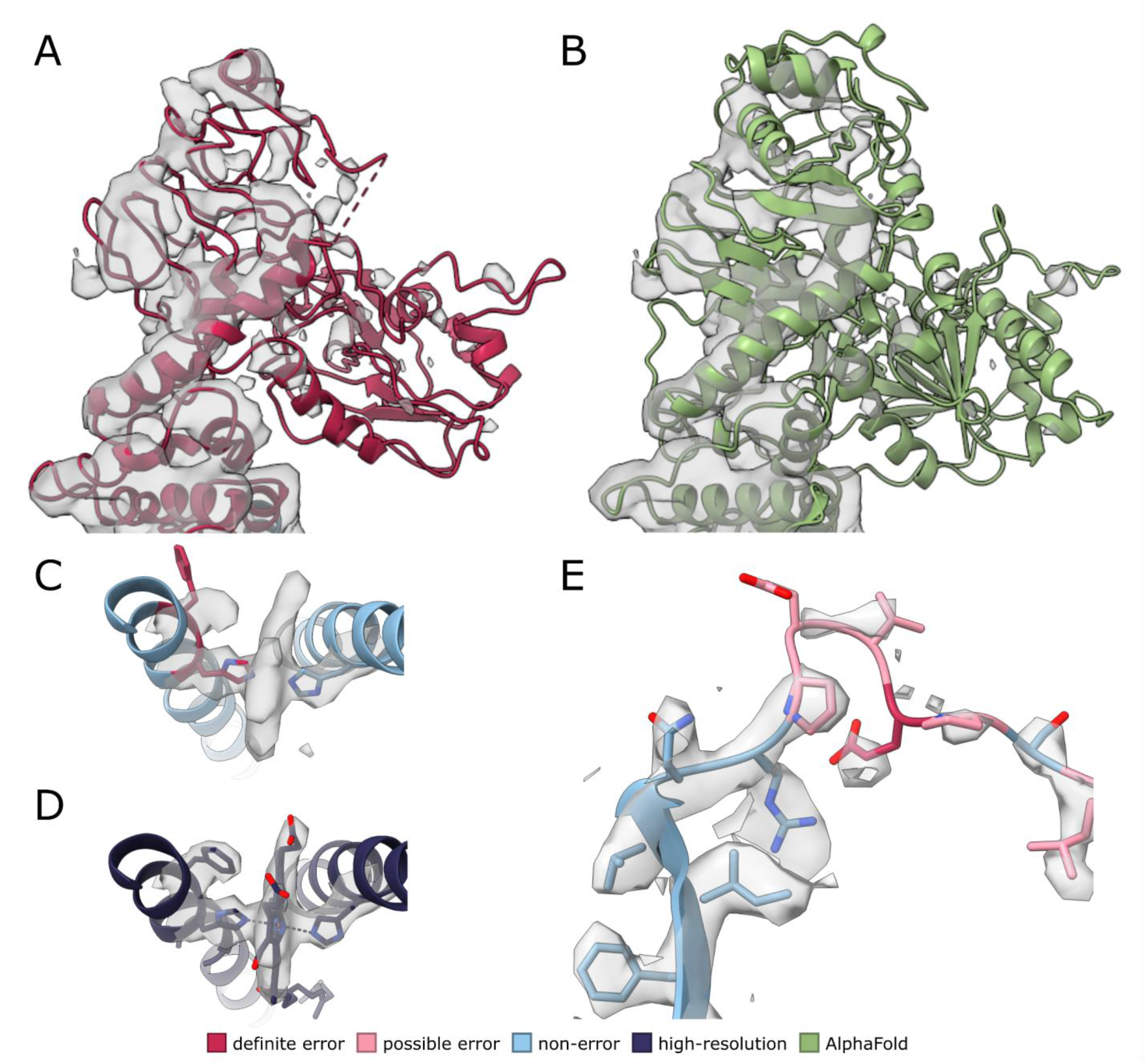
Examples of error prediction on deposited models in the EMDB. **(A)** Domain from L-fucose-1-P guanylyltransferase (PDB 5YYS) colored by MEDIC prediction. **(B)** AlphaFold prediction for protein from **A** docked into the density map. **(C)** Binding residues of cytochrome C oxidase (PDB 5Z62) after refinement and colored by MEDIC prediction. **(D)** Deposited structure of cytochrome C oxidase with ligand bound. **(E)** Rubisco activase complex (PDB 5NV3) colored by MEDIC prediction.

